# scRAPID-web: a web server for predicting protein-RNA interactions from single-cell transcriptomics

**DOI:** 10.1101/2025.03.12.642785

**Authors:** Jonathan Fiorentino, Alexandros Armaos, Chiara Montrone, Alessio Colantoni, Gian Gaetano Tartaglia

## Abstract

**Summary:** Single-cell RNA sequencing (scRNA-seq) enables high-resolution studies of gene regulation, capturing gene expression at the individual cell level. We previously developed scRAPID, a computational pipeline for predicting protein-RNA interactions and identifying hub RNA-binding proteins (RBP) and RNAs through the integration of gene regulatory network (GRNs) inference from scRNA-seq data and *cat*RAPID predictions. To make this tool accessible to a broader audience, we introduce scRAPID-web, a user-friendly web server supporting analysis of scRNA-seq data across eight model organisms. scRAPID-web offers customizable options to preprocess the input gene expression matrix, such as gene selection and cell type filtering. Users can choose from three GRN inference algorithms and decide whether to focus the analysis on specific gene types. Precompiled libraries allow fast filtering and motif-based validation of the inferred interactions. Results include detailed tables of predicted protein-RNA pairs and hubs, along with an interactive network visualization of potential RBP complexes built based on the inferred shared targets. scRAPID-web democratizes access to GRN-based analyses, providing insights into protein-RNA interactions and regulatory complexes in diverse cellular contexts.

**Availability and implementation:** scRAPID-web can be accessed at: https://tools.tartaglialab.com/scrapid.

## INTRODUCTION

Unlike bulk RNA sequencing, single-cell transcriptomics generates multiple gene expression measurements for each gene from a single biological sample, providing higher resolution for inferring gene regulatory events (Pratapa *et al*., 2020; Akers and Murali, 2021). We previously developed scRAPID (Fiorentino *et al*., 2024), a pipeline for predicting protein-RNA interactions from single-cell RNA-sequencing (scRNA-seq) data. This approach uses *cat*RAPID (Bellucci *et al*., 2011; Colantoni *et al*., 2020), a tool that predicts the interaction propensity of protein-RNA pairs based on their secondary structure and physicochemical properties, to filter the interactions returned by gene regulatory network (GRN) prediction algorithms. Among the benchmarked algorithms, DeepSEM (Shu *et al*., 2021) and TENET (Kim *et al*., 2021) emerged as the top performing in predicting protein-RNA interactions from scRNA-seq data (Fiorentino *et al*., 2024). In addition to predicting binary interactions, our approach extends to the identification of hub RBPs and RNAs (Stock *et al*., 2024). We also demonstrated that RBP–RBP interactions can be inferred by leveraging common RNA targets in predicted GRNs, providing insights into the cooperative roles RBPs play within regulatory complexes. We found that the GRNBOOST2 algorithm (Moerman *et al*., 2018) has the best performance for this task. The scRAPID toolset (Fiorentino and Armaos, 2023), available at https://github.com/tartaglialabIIT/scRAPID, is currently provided as a collection of Bash and Python scripts, making it accessible primarily to users with computational expertise. Since the pipeline can work with GRNs predicted by any tool, users are expected to run their chosen GRN inference algorithm in advance. Here, we introduce scRAPID-web, an intuitive web server that enables users to launch the scRAPID pipeline directly from a count matrix. The user can select the gene expression threshold and the gene biotypes to be retained, as well as the number of highly variable genes and the cell types to consider. Once these parameters are defined, the selected genes are used to run the chosen GRN inference algorithm. The predicted network is then filtered based on pre-computed *cat*RAPID scores and returned to the user, along with information about identified hub RBPs and RNAs. Finally, an interactive RBP-RBP network is displayed, which helps the user in retrieving putative RBP complexes.

## Description of the web server

### Precompiled libraries

scRAPID-web can be used with scRNA-seq data relative to the following organisms: *Homo sapiens, Mus musculus, Rattus norvegicus, Xenopus tropicalis, Danio rerio, Drosophila melanogaster, Caenorhabditis elegans* and *Arabidopsis Thaliana*. For each of them, we compiled RBP and RNA sequence libraries, described in **Table 1**, and we computed *cat*RAPID interaction propensities for all the possible RBP-RNA pairs. Human and mouse RBP libraries were derived from the SQL database featured in the scRAPID software. In brief, those libraries were obtained by expanding the RBP libraries from the *cat*RAPID *omics v2.0* web server (Armaos *et al*., 2021) using orthology information and integrating them with proteins scoring ≥ 10 in the RBP2GO database (Caudron-Herger *et al*., 2021). For the remaining organisms, RBPs were taken directly from the *cat*RAPID *omics v2.0* libraries and supplemented with proteins with RBP2GO score >= 10, except for *Rattus norvegicus* and *Xenopus tropicalis*, which are not covered by the RBP2GO database. Protein sequences were retrieved from Uniprot (release 2024_03) (The Uniprot Consortium, 2023).

RNA libraries were obtained from Ensembl 107 (Cunningham *et al*., 2021), except for those relative to *Arabidopsis thaliana*, gathered from Ensembl Plants 56 (Yates *et al*., 2022). Transcripts were divided into 4 categories, based on the gene biotype:

1. protein-coding: *protein_coding*;
2. long non-coding: *lincRNA, lncRNA, antisense, antisense_RNA, sense_intronic, sense_overlapping, processed_transcript, TEC*;
3. small non-coding: *ribozyme, snRNA, sRNA, snoRNA, miRNA, pre-miRNA, scaRNA, misc_RNA, vault_RNA, scRNA, Y_RNA, piRNA*;
4. pseudogene: *pseudogene, polymorphic_pseudogene, processed_pseudogene, unprocessed_pseudogene, unitary_pseudogene, transcribed_processed_pseudogene, transcribed_unprocessed_pseudogene, transcribed_unitary_pseudogene, translated_processed_pseudogene, translated_unprocessed_pseudogene, translated_unitary_pseudogene, nontranslating_CDS*.

For *Drosophila melanogaster*, transcripts at least 200 nucleotides long, with the *ncRNA* gene biotype, and containing the term “lncRNA”, “asRNA”, “sisRNA”, “Su(Ste)”, “Uhg”, “hpRNA”, or “RNaseMRP” in the gene name were classified as long non-coding RNAs. All other *ncRNAs* were included in the small non-coding RNA library.

If a gene had multiple transcript isoforms, we retained only the canonical isoform according to Ensembl annotation. When this annotation was unavailable, we chose the longest isoform. For genes with *protein-coding* biotype, we ensured that the selected transcript had a *protein-coding* transcript biotype. Transcripts are internally referenced using the *Ensembl Gene ID*.

Transcript sequences were scanned for RNA-binding motif occurrences using the same database and criteria used in the *cat*RAPID *omics v2.0* web server. For RNA-binding proteins without known motifs, we assigned motifs from the most similar RBPs with at least 70% sequence identity, if available; similar sequences were identified using MMseqs2 (Steinegger and Söding, 2017). This approach allowed us to create a dataset of RBP-RNA pairs where each RNA hosts one or more sequence motifs putatively recognized by the RBP.

For the computation of *cat*RAPID interaction propensities, we employed the fragmentation-based approach used by the *cat*RAPID fragments module (Cirillo *et al*., 2013), the *cat*RAPID *omics v2* web server (Armaos *et al*., 2021) and the RNact database (Lang *et al*., 2019). Following the approach used for RNAct and for the internal database of scRAPID, the interaction propensity for a protein-RNA pair is defined as the maximum value from the distribution of interaction propensities of their fragments. We used the sqlite3 Python package to build a SQL database of protein-RNA interaction propensities for each organism, which is fastly queried in the *cat*RAPID-based filtering step of the interactions of the scRAPID pipeline.

### Input requirements

scRAPID-web requires the following user-provided input files:

- A **gene expression matrix** from scRNA-seq or single-nucleus RNA-Seq (snRNA-seq) experiments, containing Unique Molecular Identifiers or read counts. The matrix should be in *CSV* format (compression is recommended), with genes in rows, cells in columns, gene identifiers in the first column, and cell identifiers in the first row;
- A **cell metadata file** in *CSV* format (optional unless the chosen algorithm is TENET), with cell identifiers (e.g., barcodes) in the first column (matching those in the gene expression matrix) and metadata such as cell type annotations, time points (for differentiation experiments), or any other categorization relevant for the analysis.

In addition to *Ensembl Gene IDs*, scRAPID-web also accepts gene identifiers from various categories, including *Gene Name, NCBI gene/Entrezgene accession, NCBI gene/Entrezgene ID, GenBank ID, Xenbase ID* and *ZFIN ID*. These identifiers are automatically mapped to *Ensembl Gene IDs* during the analysis.

### Analysis options

Users can configure the following parameters:

1. Organism: specify the organism from which the gene expression matrix is derived (default: *Homo sapiens*);
2. Pre-processing parameters:
  2.1 Minimum sum of counts required for a gene to pass filtering (default: 10);
  2.2 Minimum number of cells in which a gene has at least one count to pass filtering (default: 10);
  2.3 Gene types to be used used for GRN inference and downstream analyses, including messenger RNAs (mRNA), long non-coding RNAs (lncRNA), small non-coding RNAs (sncRNA) and pseudogenes (default: all).
  2.4 Number of top Highly Variable Genes (HVGs) to be selected (default: 1000). The list of genes used for the inference is the union of these HVGs with all the highly variable RBPs;
3. Gene regulatory network inference method: the available options, based on the benchmarking results obtained in (Fiorentino *et al*., 2024), are DeepSEM, TENET and GRNBOOST2 (default: DeepSEM). We suggest using TENET or DeepSEM for predicting protein-RNA interactions, while GRNBOOST2 (with 3000 HVGs) should be preferred for the prediction of RBP-RBP interactions. These are not computed when TENET is selected due to its poor performance in this task.
4. Cell metadata filtering: the user can indicate a column of the cell metadata file that contains information about cell categories (i.e. cell types or time points of differentiation) and select one or multiple options to subset the gene expression matrix. The scRAPID pipeline will be executed only on the selected categories. If TENET is chosen, the user can select an additional metadata column for the computation of diffusion pseudotime.
5. Job information: The user can indicate an email address to receive a link with the results and assign a title to the submission.

### Pipeline overview

Based on the user-specified options, the first step of the pipeline uses the Scanpy Python package (Wolf *et al*., 2018) to pre-process the gene expression matrix. This includes gene filtering (using the “scanpy.pp.filter_genes” function), normalization of the gene expression matrix (“scanpy.pp.normalize_total”), mapping of gene identifiers to *Ensembl gene IDs* and gene selection. The selection step involves applying the “scanpy.pp.highly_variable_genes” function setting the “n_top_genes” parameter according to the user-specified selection, which retains the genes with the highest variance-to-mean ratio, and ensuring that all the RBP-encoding genes classified as highly variable are included.

The available GRN inference algorithms are GRNBOOST2 (Moerman *et al*., 2018), TENET (Kim *et al*., 2021) and DeepSEM (Shu *et al*., 2021). Since each algorithm has different dependencies, we use Docker containers to ease algorithm portability. We follow BEELINE’s guidelines (https://murali-group.github.io/Beeline/BEELINE.html#adding-a-new-grn-inference-method) to ensure uniformity of input and output formats of the GRN inference algorithms, particularly for TENET and DeepSEM that were not included in the original BEELINE implementation (Pratapa *et al*., 2020). The GRN inference algorithms are integrated in the scRAPID pipeline via custom Python scripts.

TENET is the only algorithm that requires the computation of a pseudotime variable prior to GRN inference. If TENET is selected, a diffusion map (Haghverdi *et al*., 2015) is computed and an interactive plot of the first two diffusion components is displayed, with cells coloured by the categories in the selected cell metadata column. Users are requested to select a root cell for diffusion pseudotime calculation (Haghverdi *et al*., 2016) by clicking on the desired cell in the plot. A pop-up message appears to confirm the selection. To limit TENET’s runtime, if the count matrix contains more than 500 cells, a random subset of 500 cells is selected, as outlined in the original TENET implementation. After completing the preprocessing phase, the selected GRN inference method is applied, and the resulting network is filtered to include only directed RBP-target interactions. Indirect interactions are removed using precomputed *cat*RAPID interaction propensities. Hub RBPs, RNAs, and lncRNAs (if present) are identified as described in (Fiorentino *et al*., 2024). RBP-RBP interactions are scored based on the target overlap of RBP pairs, measured using the Jaccard coefficient.

### Output

The output page contains the following downloadable, searchable, and sortable tables (**Figure 1A**):

**Figure 1.**
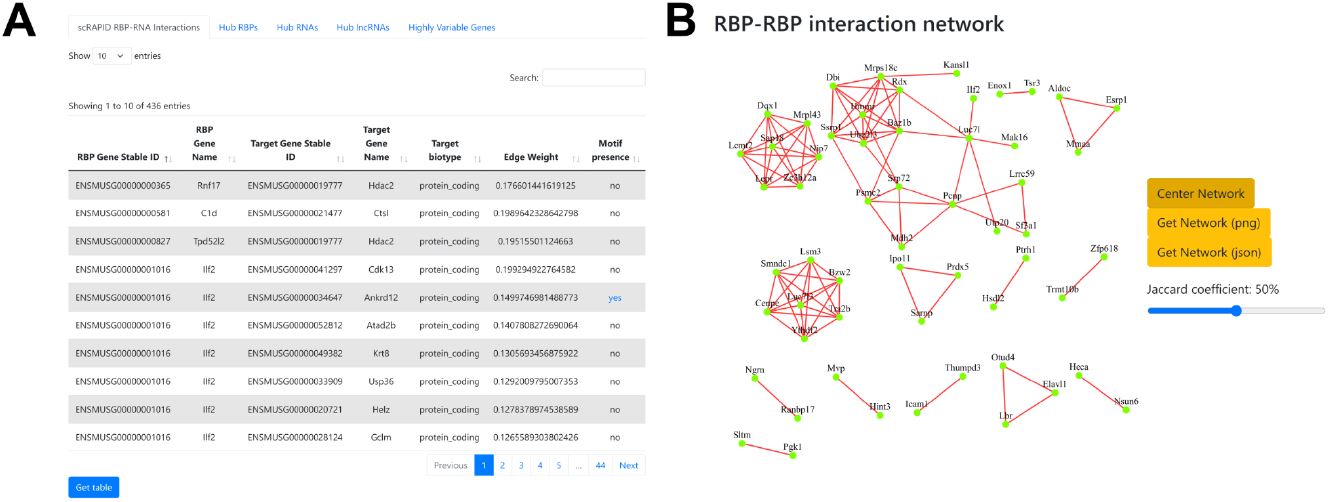
scRAPID-web output page. (**A**) Table reporting the protein-RNA pairs inferred by the selected GRN inference method and predicted to interact by *cat*RAPID. Additional output tables can be accessed by clicking on the corresponding tabs. (**B**) Interactive RBP co-interaction network, where users can drag and reposition nodes (RBPs) and adjust the Jaccard distance cutoff to filter displayed edges (predicted RBP-RBP interactions).

1. scRAPID protein-RNA interactions: list of inferred protein-RNA interactions that passed the *cat*RAPID-based filter, including target RNA biotype, edge weights returned by the selected GRN inference method and RBP-specific motifs identified within the target sequences;
2. Hub RBPs: list of RBPs and their out-centrality values (fraction of outgoing edges in the GRN);
3. Hub RNAs: list of target RNAs and their in-centrality values;
4. Hub lncRNAs: list of target lncRNAs, if present in the dataset, including in-centrality values (fraction of ingoing edges in the GRN);
5. RBP co-interactions: list of RBP-RBP pairs including the number of target RNAs of both RBPs from the inferred GRN and the Jaccard coefficient quantifying target overlap.
6. Highly Variable Genes: list of the highly variable genes on which the inference is run.

At the top of the output page, an interactive RBP-RBP interaction network is displayed, where nodes represent RBPs and edges indicate interactions with Jaccard coefficient exceeding a user-defined threshold (**Figure 1B**). Users can adjust the threshold dynamically using a slider, which automatically updates the network layout in real time. The network can be downloaded as a *PNG* image or a Cytoscape-compatible *JSON* file.

### Web server implementation

scRAPID-web features a Python Flask backend, implemented as a RESTful API.The queue system ensures efficient processing of multiple user requests by leveraging Celery workers and Redis for task management. The front-end interface is designed using JavaScript, HTML, and styled with Bootstrap and jQuery, incorporating Font Awesome for enhanced visual elements. The frontend communicates dynamically with the backend to ensure a smooth Single-Page Application experience. The network visualization is implemented using Cytoscape.js, enabling an interactive and dynamic user experience.

## CONCLUSIONS

We developed a web server enabling users to run the scRAPID pipeline on scRNA-seq data from eight different organisms, significantly expanding the range of supported organisms beyond those included in the internal libraries of the standalone version of scRAPID. Additionally, the inferred RBP-RNA interactions which are supported by RNA-binding motif occurrences are reported, offering an orthogonal layer of validation. The web server features a user-friendly graphical interface, enabling non-expert users to seamlessly utilize advanced GRN inference algorithms that typically require GPU configuration and multiple dependencies. Finally, the interactive RBP-RBP network offers an intuitive platform to explore potential protein complexes formed in specific cellular contexts.

## ACKNOWLEDGMENTS

The authors would like to thank the ‘RNA initiative’ at IIT and all the members of Tartaglia’s lab at CRG, Sapienza and IIT.

## Author contributions

A.A. and J.F. developed the web server. J.F. built the computational framework with the aid of C.M. C.M. tested the web server and wrote documentation and tutorial. A.C. built the precompiled libraries. A.C. and G.G.T. conceptualized the work and wrote the manuscript, with the contribution of J.F. and A.A.

## FUNDING

The research leading to this work was supported by the ERC ASTRA_855923 (G.G.T.), EIC Pathfinder IVBM4PAP_101098989 (G.G.T.) and PNRR grant from National Centre for Gene Therapy and Drugs based on RNA Technology (CN00000041 EPNRRCN3 (G.G.T.). F.D.P. acknowledges the support granted by the European Union - Next Generation EU, Mission 4 Component 1 CUP D53D23016360001, PRIN-PNRR, Grant n. P2022CLXMK, for his current position.

## LEGENDS

**Table 1.** Precompiled RBP and RNA libraries available in scRAPID-web for each available organism. lncRNA: long non-coding RNA; sncRNA: small non-coding RNA.

## Notes

https://tools.tartaglialab.com/scrapid

### Competing Interest Statement

The authors have declared no competing interest.

